# AutoRNC: an automated modeling program for building atomic models of ribosome-nascent chain complexes

**DOI:** 10.1101/2023.06.14.544999

**Authors:** Robert T. McDonnell, Adrian H. Elcock

## Abstract

The interpretation of experimental studies of co-translational protein folding often benefits from the use of computational methods that seek to model the nascent chain and its interactions with the ribosome. Ribosome-nascent chain (RNC) constructs studied experimentally can vary significantly in size and the extent to which they contain secondary and tertiary structure, and building realistic 3D models of them therefore often requires expert knowledge. To circumvent this issue, we describe here AutoRNC, an automated modeling program capable of constructing large numbers of plausible atomic models of RNCs within minutes. AutoRNC takes input from the user specifying any regions of the nascent chain that contain secondary or tertiary structure and attempts to build conformations compatible with those specifications – and with the constraints imposed by the ribosome – by sampling and progressively piecing together dipeptide conformations extracted from the RCSB. We first show that conformations of completely unfolded proteins built by AutoRNC in the absence of the ribosome have radii of gyration that match well with the corresponding experimental data. We then show that AutoRNC can build plausible conformations for a wide range of RNC constructs for which experimental data have already been reported. Since AutoRNC requires only modest computational resources, we anticipate that it will prove to be a useful hypothesis generator for experimental studies, for example, in providing indications of whether designed constructs are likely to be capable of folding, as well as providing useful starting points for downstream atomic or coarse-grained simulations of the conformational dynamics of RNCs.

## Introduction

In recent years there has been considerable interest in the development and application of methods for studying co-translational protein folding (for recent reviews, see (1–4)). While the motivations for such studies vary, they typically involve the generation of ribosome-nascent chain complexes (RNCs) of constructs designed specifically to provide insights into the ability of proteins to fold. Impressive advances in experimental techniques, in particular, have facilitated probing the extent of co-translational protein folding that can occur as nascent chains: (a) work their way through the ribosome exit tunnel (e.g. (5–12)), (b) interact with the ribosome surface (e.g. (13–19)), and (c) interact with proteins being synthesized by other ribosomes (20). Many of these experimental studies have been used in combination with computational methods that are intended to statically model and/or dynamically simulate the RNC constructs (e.g. (8,10–12,14,20–22)). Molecular simulations of co-translational folding events with detailed ribosome models date back many years (23), but while simulation-focused studies of co-translational folding have provided a number of interesting insights (e.g. (23–25)), it is more usual these days for computational studies to be performed in combination with experimental studies of the same constructs: in this way, most of the advantages of the computational methods can be enjoyed while the experimental data provide key restraints that keep the simulations “honest”.

One difficulty in applying computational methods to the modeling of RNCs is the paucity of available methods to facilitate the building of plausible conformations of the nascent chain on the ribosome. Recent years have seen dramatic advances in the ability of computational methods to accurately predict the atomic structures of folded proteins (26), with the development of AlphaFold2 (27) being a clear watershed moment for computational structural biology. In their present forms, however, such methods do not make predictions of proteins subject to the constraints that would be imposed by the ribosome tunnel. For this reason, we have developed an automated computational method that we call AutoRNC that attempts to build RNCs subject to constraints specified by the user. Full details of the AutoRNC methodology are provided in Methods; the remainder of this manuscript describes potential use-cases for the code, focusing on RNC constructs that have been studied experimentally.

## Results

A schematic illustration of the AutoRNC method is provided in Figure 1. The process can be briefly outlined as follows. The user provides the sequence of their nascent chain construct and selects one of the ribosome template structures provided with the code (or provides a new one) into which the construct is to be built. If desired, the user can also specify details of any secondary or tertiary structure elements that are known (or hypothesized) to be present in the construct. If tertiary structure elements are defined, then the user also provides a full-length atomic structure of the construct. In all the cases containing tertiary structure that are presented here, these atomic models were predicted using the ColabFold derivative (28) of AlphaFold2 (27), but experimental or homology modeled structures are equally acceptable provided that they do not contain any stretches of missing residues. Given this information, together with additional specified input parameters, AutoRNC then attempts to generate models by building the nascent chain vectorially starting from the construct’s C-terminal residue, which is usually assumed to be directly attached to the peptidyl transfer center (PTC) tRNA of the ribosome template structure. AutoRNC builds the nascent chain by piecing together dipeptide conformations that are either sampled from libraries that we have constructed from the RCSB ((29); see Methods) or that are taken directly from the atomic structure provided by the user (see above). If a given dipeptide superimposes poorly on to the previously added residue or if the newly added residue experiences a steric clash (either with other, pre-placed atoms of the nascent chain or with atoms of the ribosome), then the proposed dipeptide conformation is rejected. Depending on the situation (see Methods), the code then either: (a) randomly selects a different conformation of the dipeptide, (b) backtracks by a user-selected number of residues (N_back_), or (c) rejects the partially built chain entirely and restarts from the C-terminal residue of the construct. The code continues in this way until all residues of the nascent chain have been placed; it then repeats the entire process until the desired number of models has been built. If the user’s construct has specified features that are structurally compatible with the steric constraints imposed by the ribosome’s exit tunnel (see below), AutoRNC will typically build hundreds of plausible models of the nascent chain in a matter of minutes.

**Figure 1.**
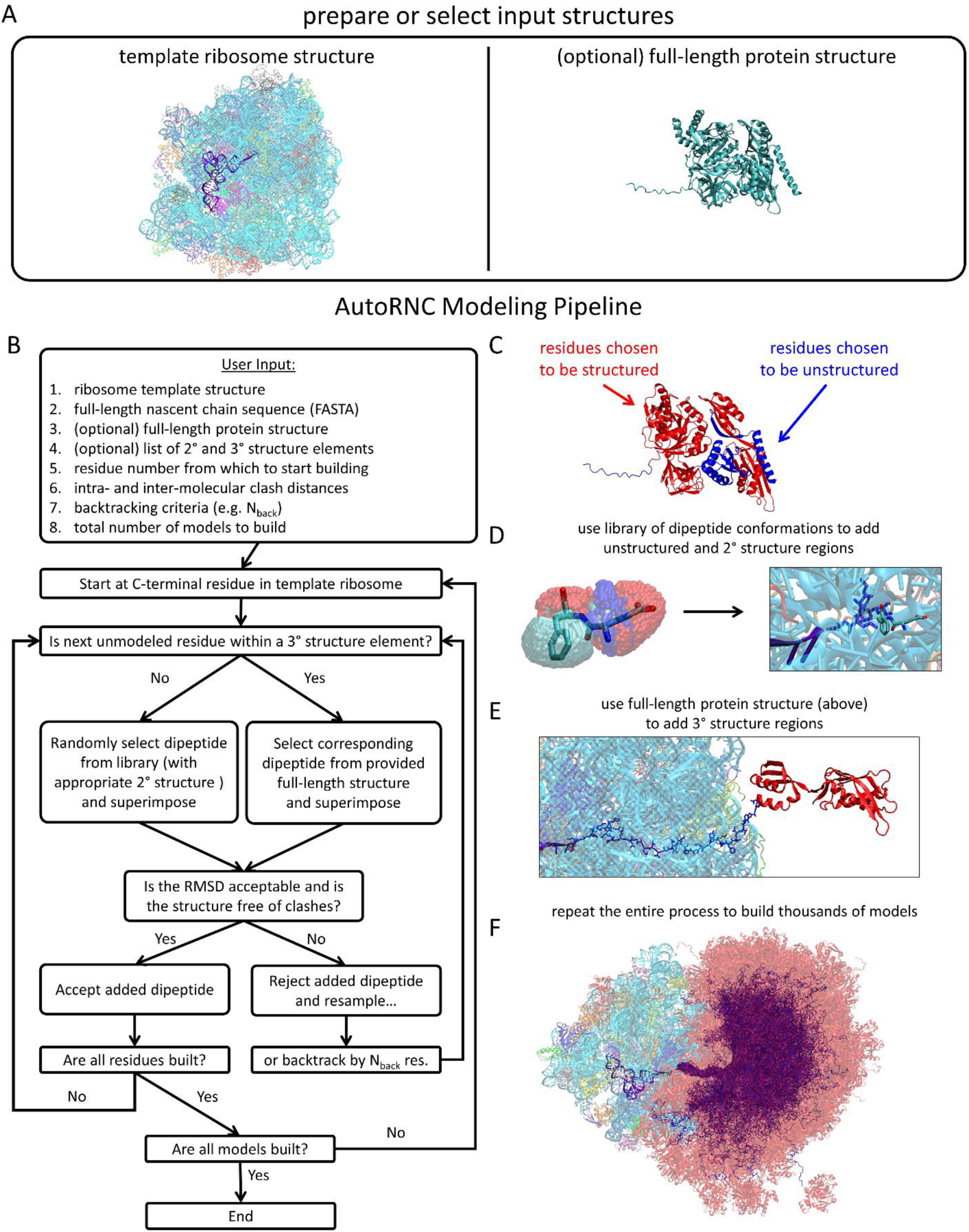
Schematic illustration of the AutoRNC modeling pipeline. (A) AutoRNC requires that the user selects (or alternatively, provides) a ribosome template structure (transparent cartoon) containing a P-site tRNA (opaque purple cartoon) from which to build the desired nascent chain. To model tertiary structure, users optionally can provide a folded structure of their protein of interest. (B) Flow chart of the AutoRNC modeling pipeline. (C) Users can exert control over which residues to model as unstructured, and which to model as members of 2° or 3° structure elements. AutoRNC builds the nascent chain from the C-terminal residue using: (D) dipeptide conformation libraries for unfolded or secondary structured regions, or (E) an input protein structure for tertiary structured regions. (F) The entire modeling process repeats until the desired number of models has been built; thousands of models can be built in a single run.

While AutoRNC can build plausible models of constructs that contain folded or unfolded domains, its predictive abilities are essentially restricted to modeling: (a) the conformational properties of unfolded or disordered stretches within the nascent chain, and (b) the ways in which such stretches, and any structured elements specified by the user, can be placed subject to the constraints imposed by the ribosome. Two validations of AutoRNC’s capabilities in the first of these roles are provided in Figure S1. In Figure S1A we show histograms of the radius of gyration, R_gyr_, of four proteins whose unfolded states have been characterized experimentally by the Clark, Plaxco and Sosnick groups (reviewed in (30)). For these calculations, we used AutoRNC to build 10,000 models of each protein in the absence of any ribosome template. In all four cases, the mean R_gyr_ values obtained with AutoRNC using its default input parameters agree well with the experimental values. For protein L (64 amino acids), ubiquitin (76 AA), acylphosphatase (98 AA), and the intrinsically disordered N-terminal domain of pertactin (334 AA), we obtain mean R_gyr_ values of 22.7, 25.8, 29.7, and 55.9 Å respectively, which are to be compared with the corresponding experimental estimates of 25 (31,32), 25 (33), 30 (33), and 60 Å (34). In Figure S2B, we show the Ramachandran angle distribution computed from 1000 models generated by AutoRNC of a randomized 1000-residue sequence, again in the absence of any ribosome. Given that AutoRNC builds fully unfolded chains by sampling dipeptides extracted from coil regions of the RCSB (see Methods), it is perhaps not surprising that the distribution of Ramachandran angles in the generated conformations appears as it does: all of the expected regions of the Ramachandran map are visited although for the particular sequence modeled here, there appears to be a slight preference given to α-helical conformations.

While the above results demonstrate that AutoRNC can build realistic models of proteins in their unfolded states, the code’s primary purpose is to build models of RNC constructs that are sufficiently plausible that they can be used to either interpret existing experimental data or to pre-screen candidate constructs for subsequent experimental study. In the Figures that follow, we demonstrate AutoRNC’s use in a variety of scenarios. In cases where the available experimental data indicate that the nascent chain remains essentially unfolded within the ribosome tunnel, we have used AutoRNC to build complete atomic models; in those cases where the data indicate that secondary and/or tertiary structure forms within the tunnel, we have opted to accelerate the AutoRNC calculations by building nascent chain models that contain complete backbone coordinates but omit the sidechains (see Methods). In all the examples shown, we focus our attention on AutoRNC models in which the nascent chain follows the generally accepted path of the ribosome tunnel. Toward the end of the manuscript, however, we show that AutoRNC’s models can also sample alternative exit tunnels and that they can sometimes reflect other idiosyncrasies of the ribosome templates used in model generation.

To illustrate AutoRNC’s use as a method for visualizing and interpreting known experimental data, Figure 2 shows example models produced for a range of constructs described in the literature; these illustrated examples cover all the major use cases of the code. Figure 2A shows AutoRNC’s use on an RNC construct that is thought to be entirely unfolded; for such cases, the user needs to provide only the sequence. The particular construct shown is one studied by the Bustamante group: it is 204 residues in length and includes the first 167 residues of *S. erythraea* calerythrin (35); details of this and all other constructs presented in this work can be found in Table S1. The Figure shows an overlay of 1000 such models and gives a sense of the ability of the nascent, unfolded chain to sample conformations that stretch a long distance from the exit of the ribosome tunnel.

**Figure 2.**
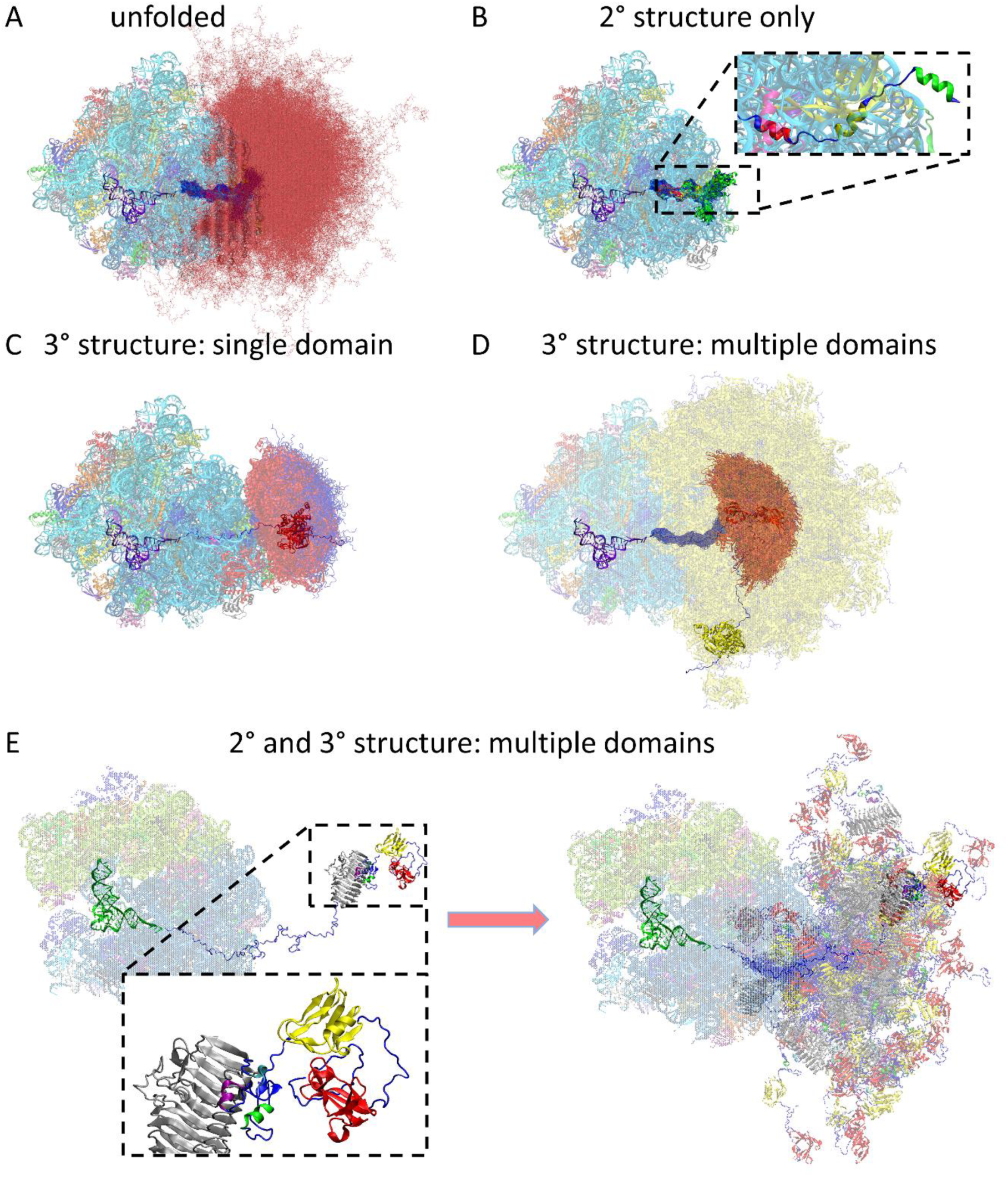
AutoRNC can model constructs with arbitrary combinations of structural elements. Each panel highlights nascent chain conformations with different structured statuses modeled by AutoRNC. (A) Unfolded nascent chain models of a 204-residue calerythrin construct (35); blue lines represent the C-terminal residues 128-167, red lines represent residues 1-127). (B) Secondary structure in nascent chain models of a 56-residue HemK construct (5,7) all contained within the ribosome exit tunnel; the insert highlights a single conformation with helices colored red, yellow and green. (C) Nascent chain models with a single folded domain from a 478-residue EF-G construct (17); combined domains G+II (residues 16-421) are colored red, all other residues blue. (D) Nascent chain models with multiple, independently folded domains from a 730-residue EF-G construct (36); combined domains G+II (residues 16-425) are now yellow; combined domains IV+V (residues 506-695) are red; the intervening domain III (residues 431-503) and the C-terminal 35 residues are blue. (E) Nascent chain models with both secondary and tertiary structure from a 686-residue P22 tail spike protein construct (37); α-helices are shown as green, purple and cyan cartoons; tertiary elements are shown as grey, yellow and red opaque cartoons). Left image shows a single selected model; right image shows many models overlaid.

Figure 2B illustrates AutoRNC’s use on a construct known to contain elements of secondary structure only; for these cases, the user needs to supplement the sequence with a text file identifying those elements of secondary structure that are desired. The specific construct shown is one that contains the first 56 residues of the *E. coli* protein HemK studied by the Rodnina group (5,7); while apparently not capable of forming stable tertiary structure, this construct is thought to contain α-helices at residues 4-12 (green), 20-29 (yellow), and 36-41 (red). AutoRNC ensures that lists of secondary structure elements provided by the user are respected in the final models; in the specific model shown in the inset of Figure 2B, however, it can be seen that the code may also occasionally choose to extend elements of secondary structure if it happens to randomly sample appropriate dipeptide conformations. It should be noted that while the conformations in the dipeptide library used for unfolded regions were specifically extracted from coil regions of protein structures, they nevertheless contain isolated residues that populate the α-helical and β-sheet regions of the Ramachandran map (see Figure S1B).

Figures 2C and 2D show that AutoRNC’s capabilities extend beyond building constructs that contain isolated stretches of secondary structure to constructs that contain one or more folded globular domains. The two example constructs shown come from the Kaiser group’s experimental studies of the multi-domain, 704-residue *E. coli* protein EF-G (16,17,36). Figure 2C shows models of a 478-residue construct, which is sufficiently long that EF-G’s domains G and II are both thought to be natively folded. Figure 2D shows a larger 730-residue construct which is thought to contain two separate folded blocks: the first, composed of domains G & II, and the second composed of most of the discontinuous domain IV, together with all of domain V. These two folded blocks are thought to be separated by a stretch of unfolded residues that in the full-length native protein would correspond to domain III. Figures 2C and 2D show that both constructs can be modeled by AutoRNC under the assumption that the folded domains each take on the conformation seen in the structure of the full-length protein.

Finally, Figure 2E shows that AutoRNC can handle combined scenarios in which some regions of a RNC construct adopt tertiary structure while others contain only secondary structure. In this case the construct is one studied experimentally by the Clark group (37): it is 686 residues long and includes the entire sequence of bacteriophage P22’s tail spike protein. For this construct, the exact elements of secondary and tertiary structure present in the nascent chain are less well defined experimentally, but it is thought that the first two-thirds of the residues are largely folded, while the remainder are probably unfolded. Given this information, we have chosen, purely for illustrative purposes, to select those structural elements that are present in the construct’s AlphaFold2-predicted structure for residues 1-542 and have modeled the remaining residues as unfolded. Despite the length of the construct, and the complicated arrangement of secondary and tertiary structure within the nascent chain, large numbers of models can be rapidly generated by AutoRNC.

AutoRNC’s speed of execution is sufficiently great – especially when sidechains are omitted from the nascent chain or when the steric clash criteria are relaxed (see Methods) – that it often becomes feasible to generate models for all possible lengths of a given nascent protein; this in turn allows users to develop a structural and visual representation of the trajectory that a protein may take at all stages of its synthesis. Figure 3 shows three examples of AutoRNC’s use to illustrate folding trajectories that have been determined experimentally. Figure 3A shows four constructs of *E. coli* HemK studied by the Rodnina group with lengths ranging from 35 to 112 residues (5,7). The experimental data have indicated that these constructs can acquire secondary structure, in the form of α-helices, that fold within the ribosome tunnel; only in the longest construct, however, are these helices capable of maintaining the tertiary contacts necessary for them to adopt a stable fold. When sidechain atoms are omitted from the calculations (see above), AutoRNC rapidly builds models for all constructs, including the longest one in which the folded domain fits snugly into the vestibule that exists at the end of the ribosome tunnel (see far panel of Figure 3A).

**Figure 3.**
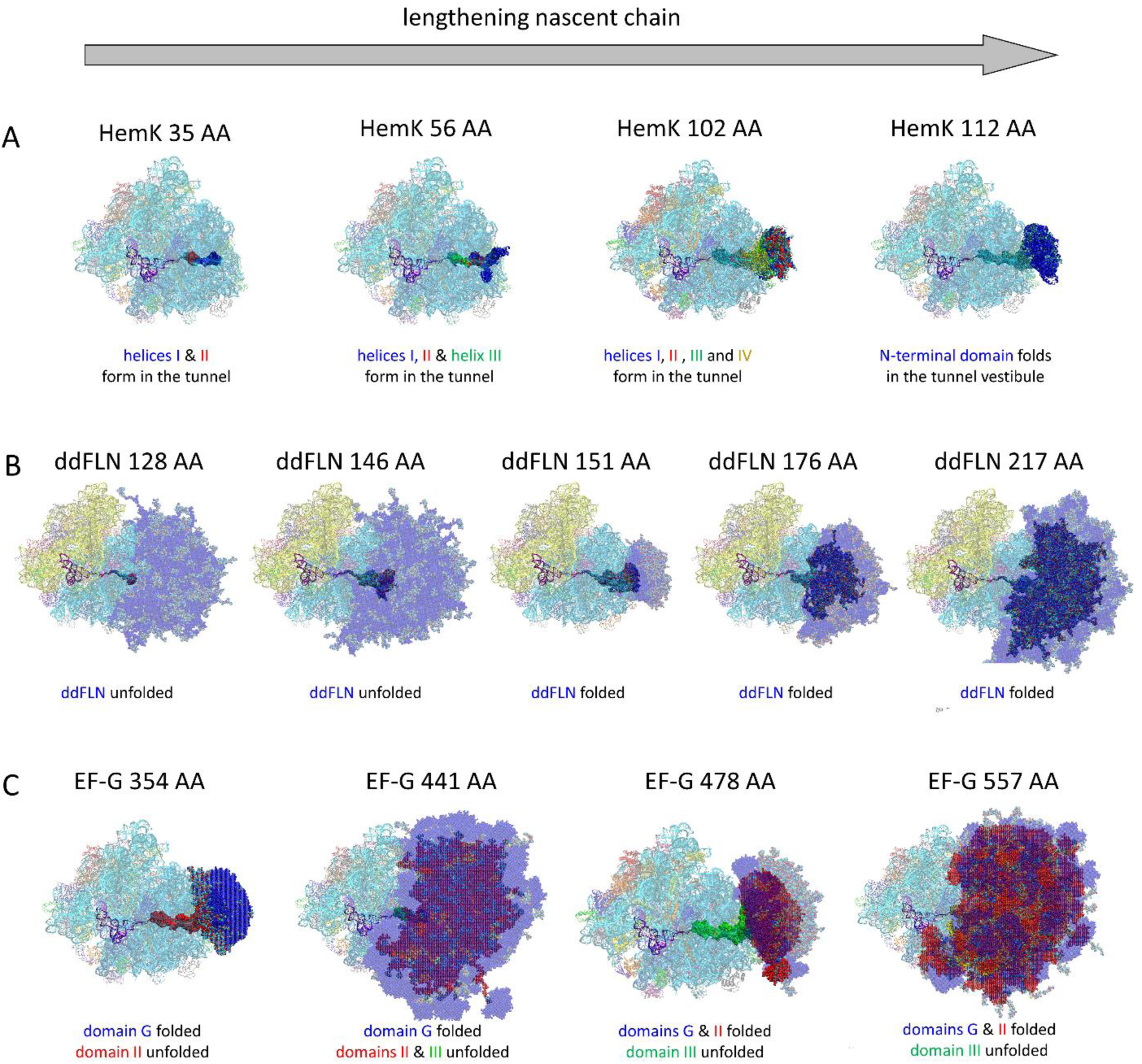
AutoRNC models can illustrate co-translational folding trajectories of RNCs. Each panel shows 1000 RNC conformations modeled by AutoRNC for constructs that fold co-translationally in experiment as the nascent chain increases in length (from left to right) and emerges from the ribosome exit tunnel. (A) *E. coli* HemK constructs (5,7) in which α-helices I (blue), II (red), III (green) and IV (yellow) form within the ribosome exit tunnel and fold into the N-terminal domain (blue, HemK 112 AA) once the nascent chain is sufficiently long. (B) *D. discoideum* Filamin-like domain constructs, “ddFLN” (39) in which the domain folds only after emerging from the ribosome exit tunnel. (C) *E. coli* EF-G constructs (17,36) in which domains G (blue), II (red) and III (green) extend out of the ribosome exit tunnel; in the constructs shown, only domains G and II are capable of folding. All models shown here were created using folded states determined experimentally and ribosome template structures consistent with the method used to stall translation experimentally.

Figure 3B shows five progressively longer constructs ranging in length from 128 to 217 residues of the *<u>D</u>. <u>d</u>iscoideum* immunoglobin-like filame<u>n</u> domain (ddFLN) studied by the Christodoulou group (38,39). In this case, the experimental data have indicated that residues 9-109 fold into their native structure only in constructs that contain 151 or more residues. The models generated by AutoRNC respect this feature but also suggest that folding of the ddFLN domain is accompanied by an abrupt change in the ability of the nascent chain to extend out into the surrounding environment. Interestingly, since it contains no stably folded structure, the smaller 1-146 construct reaches considerably further out into solution than a construct (1–176) that contains an additional 30 residues (compare second and fourth panels of Figure 3B).

Finally, Figure 3C shows a series of four constructs that map the early to middle stages of synthesis of *E. coli* EF-G studied by the Kaiser group (16,17,36). The experimental data indicate that the first two of the illustrated constructs contain a fully folded G domain (residues 16-308), while the remaining two constructs are sufficiently long that domain II (residues 310-421) also becomes fully folded. AutoRNC’s models of these constructs satisfy these constraints but again provide insights that are not immediately obvious from examining only the folding status of the individual domains. For example, while domain G is fully folded in all four constructs, its disposition relative to the ribosome oscillates substantially depending on both the length of the construct and the folding status of the residues that intervene between it and the ribosome’s PTC. In particular, echoing what was shown with ddFLN (Figure 3B), the distance between domain G and the ribosome exit tunnel decreases when the number of intervening unfolded residues decreases, even in cases in which this is accompanied by an increase in the *total* number of intervening residues.

Along with sampling the conformational possibilities open to the nascent chain once it extends beyond the ribosome tunnel, AutoRNC can also provide effective sampling of potential interactions between the nascent chain and the interior of the ribosome tunnel. In particular, the ability to rapidly generate large numbers of atomic models allows users to determine the extent to which individual residues can access different regions of the tunnel in atomic detail. Figure 4, for example, shows overlays of 1000 models of a 441-residue construct of EF-G studied by the Kaiser group, with each panel highlighting the positions sampled by a different residue. Importantly, such views are made in the context of complete conformational models of the construct of interest. This makes it possible to account explicitly for the fact that the ability of sidechains nearer the C-terminus of a construct to access certain regions within the ribosome’s tunnel might be a function of the folding status of residues that are outside the tunnel.

**Figure 4.**
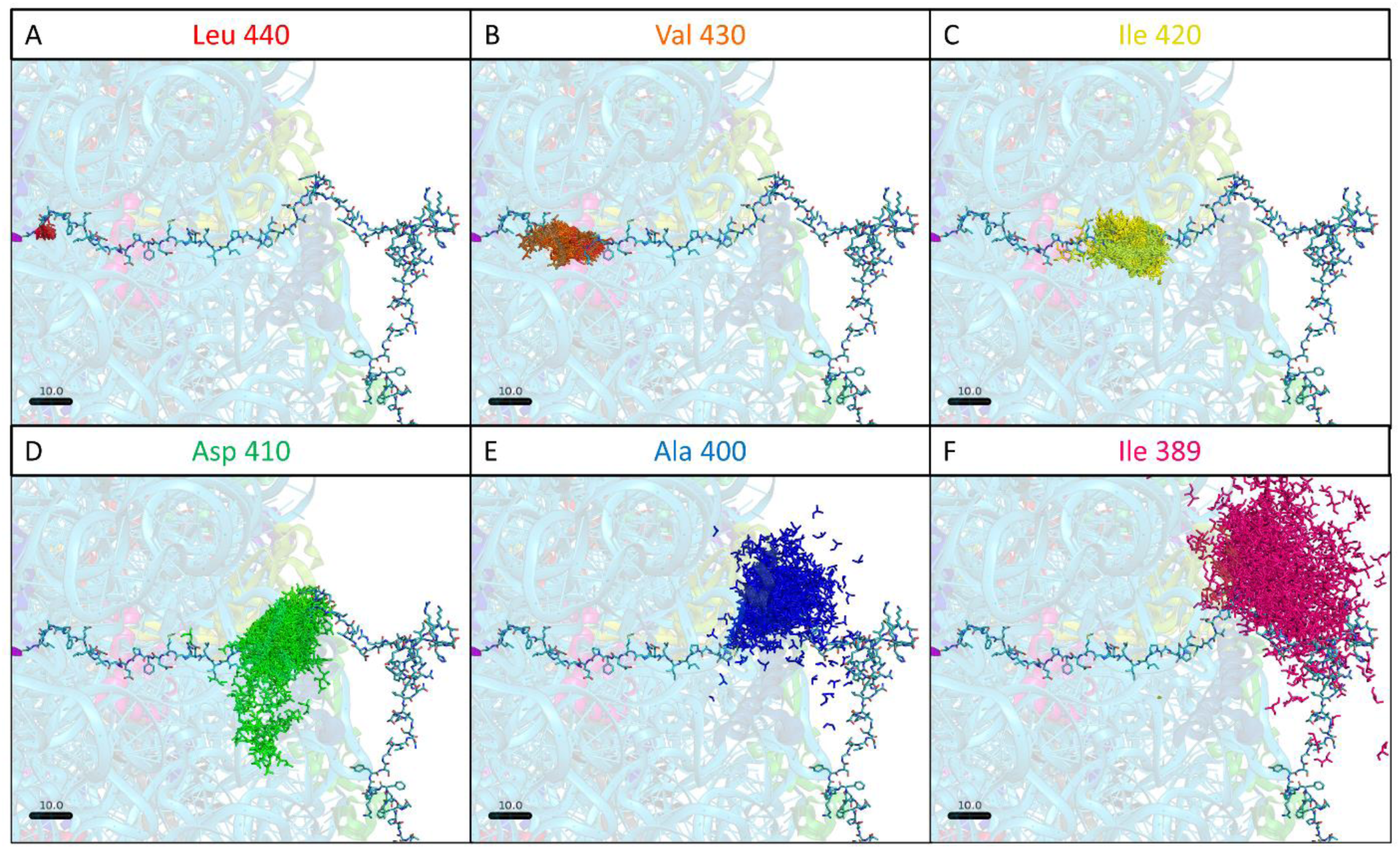
AutoRNC samples residue positions within the ribosome exit tunnel. (A-F) 1000 different models of selected nascent chain residues from a 441-residue EF-G construct (17). Each panel shows a different selected residue, displayed with a single conformation of the entire construct.

The images shown in Figures 2 to 4 provide a representative sampling of the RNC constructs that we have built with AutoRNC. In total, we have built models of 59 experimentally studied RNC constructs taken from the literature. Figure S2 shows a gallery of all modeled RNCs that use an elongating ribosome structure (see Methods); Figure S3 shows those that use ribosome templates that contain the SecM and TnaC arrest peptides. In most cases, AutoRNC rapidly built models consistent with the experimental data using its default input parameters. In a minority of cases, models were returned more slowly by AutoRNC, and this usually reflected difficulties in building sterically acceptable atomic models for folded elements within the ribosome exit tunnel. To obtain models more rapidly, we used one of two alternative strategies (see Methods), either: (a) building models that retain only the sidechain’s backbone atoms – as was done for some of the constructs shown in Figures 2 and 3 (see above) – or (b) decreasing the usual clash distance criteria to be more tolerant of steric clashes. In two cases, despite using these alternative strategies, AutoRNC still initially failed to produce any acceptable models. Interestingly, both of these problematic constructs involved folded domains with relatively short linkers connecting them to the PTC and both made use of the same ribosome template structure, i.e., the 7OIZ structure that we use for nascent chain constructs containing a TnaC arrest peptide. The cause of the difficulty appears to be a loop on ribosomal protein L24 (residues 45-56) that juts out into the middle of the ribosome exit tunnel (see Figure S4A). A similar issue with L24 has caused at least one other group to remodel this loop for simulations (11); here, we have adopted a temporary strategy of simply deleting the residues of the loop to accelerate model building.

The obstructive role played by L24 in the 7OIZ (and possibly other) ribosome structures is not the only case where an idiosyncrasy of the selected ribosomal template appears to have an impact on the models generated by AutoRNC. We can point to two more cases where the ribosome structure leads to RNC models that are of uncertain significance and that users should be aware of when using the AutoRNC code. The first and most interesting example is the finding that a sizeable minority of models that are generated by AutoRNC do not follow the canonical ribosome exit tunnel, but instead follow one or more alternative pathways (see Figure S4B). The possibility that there might be alternative exit tunnels has been raised previously in the literature (e.g. (40)); what appears to be new here is the finding that AutoRNC indicates that these pathways are sufficiently wide to physically accommodate plausible structural models of nascent chains: it is to be noted that the example shown in Figure S4B was from a fully atomic modeling run performed by AutoRNC with the default clash parameters. In this case, the nascent chains sample three distinct exit tunnels similar to those suggested many years ago by the Frank group (40): (1) the “canonical” exit tunnel that provides the shortest exit path and is located between ribosomal proteins L22 (yellow), L24 (green) and L29 (dark blue), (2) “alternative exit 1” located near L4 (dark grey) and (3) “alternative exit 2” located adjacent to L24 and L29.

A second scenario in which AutoRNC’s models occasionally follow unexpected pathways appears to us to be less plausible. Especially in cases where a nascent chain possesses features that make it difficult to find sterically acceptable conformations, we have found that the nascent chain can occasionally “double-back” on itself and occupy open spaces around the ribosome’s bound tRNAs (see Figure S4C). We leave the question of the likely significance of these findings to future work, but we note that nascent chain models that follow non-canonical pathways can be eliminated if desired by either post-processing of the models or by artificially blockading the undesired pathway(s) with added atoms that prevent AutoRNC from successfully inserting a nascent chain.

## Discussion

We think that AutoRNC’s ability to rapidly build plausible atomic models of RNCs means that it may be of use in a number of different scenarios. One obvious use is to generate conformational landscapes for RNC constructs that have already been studied experimentally; its capabilities in this scenario can be gauged by noting the ∼60 different constructs from the literature that AutoRNC has already been used to model (see Supporting Information). A second potential use of the code is to guide the selection of constructs prior to experimentation: constructs that are hypothesized to be capable of folding but for which AutoRNC has difficulty generating atomic models may in some cases not be capable of folding in the way imagined (for exceptions to this, see below). A third straightforward use for AutoRNC is to generate starting configurations for computer simulation studies that use molecular dynamics or other techniques to model the conformational dynamics of nascent chains on the ribosome.

In considering the potential applications of AutoRNC, it is important to be clear about what the code does and does not do. AutoRNC does not explicitly predict the degree of secondary structure present in a nascent chain construct, nor does it explicitly predict the tertiary folding status of individual domains of a nascent chain. These are things that the user must either know, or must frame hypotheses about, prior to using the code. On this basis, therefore, it might be argued that much of what appears in the models that are generated by AutoRNC is simply what was asked for by the user. There is clearly some truth to such a view, and it seems reasonable to imagine that many potential users may wish to use AutoRNC simply to make attractive figures that illustrate, in explicitly structural terms, data and insights that have largely been arrived at by direct experimentation.

We also think, however, that the atomic models produced by AutoRNC provide additional information that can only be obtained by building detailed structural models of RNCs. Some examples of phenomena that we think can be examined using AutoRNC’s models include: (a) changes in the degree to which nascent chains extend into the environment as their domains fold or their inter-domain linkers lengthen, (b) changes in the ribosome surfaces that are contacted by folded domains when the remainder of the nascent chain either folds or lengthens, and (c) changes in the physicochemical environments that individual amino acid sidechains experience in the ribosome tunnel as the nascent chain lengthens. Importantly, all of these aspects can be examined in the context of complete conformational models of the construct of interest.

While AutoRNC does not explicitly predict the folding status of the domains within a given construct (see above), it seems likely that in the longer term AutoRNC may be able to determine the extent to which confinement within the ribosome tunnel might limit an individual domain’s ability to fold. In particular, the relative speed with which AutoRNC produces models of a series of candidate constructs might provide a rough indication of the extent to which they are likely to be folded in an experimental setting. It may be possible to use the current version of the code in this way already, but it should be remembered that in application to ∼60 experimentally studied constructs we encountered several cases where AutoRNC’s criteria for generating models had to be relaxed in some way, and two cases where the ribosome’s L24 loop had to be eliminated, for complete models to be returned. The need to occasionally relax the otherwise stringent criteria applied during AutoRNC’s model-building efforts almost certainly results from using single, static ribosome templates. A longer-term strategy might be to allow the ribosome to “breathe” by including a range of ribosome conformations in the calculations. If such an extended version of AutoRNC can be shown to rapidly build all known experimental RNC constructs using a single protocol, without the need to adjust parameters on a case-by-case basis, then it could be a reliable method for pre-screening hypothetical constructs for viability in advance of their experimental study.

Finally, we note a few other directions for applications or future developments of the code. One immediate area of application – i.e., one that requires no modification to AutoRNC’s source code – is to the modeling of nascent chains in eukaryotic ribosomes: the AutoRNC code is entirely agnostic regarding the origins of the ribosome structures that are used in the calculations. The release version of AutoRNC contains three ribosome template structures that are ready for use with the code; reflecting the constructs typically studied by the experimental community, however, all such structures are of the *E. coli* ribosome. Additional template structures will be added in future updates of the code – e.g., chaperone-bound structures of the *E. coli* ribosome – but since the steps needed to prepare structures for use are relatively straightforward, it is possible for users to develop their own if required.

A second direction for future work is to develop methods that allow conformations with desired properties to be more easily sampled. Currently, conformations are accepted or rejected by AutoRNC solely according to how well the sampled dipeptide conformations superimpose on each other and whether they are free of steric clashes between the nascent chain and the ribosome. It would obviously be helpful, however, to be able to up-weight conformations that are likely to have especially favorable energetic interactions. Accounting for such features during AutoRNC’s sampling of chain conformations is not likely to be easy to implement, so alternative approaches are probably worth pursuing first. In principle, the simplest way to identify more favorable nascent chain conformations is to subject AutoRNC’s models to molecular dynamics simulations. A downside to such an approach is obviously the considerable computational expense of performing such simulations, so an alternative way that seems worth exploring is to add a post-processing step that reweights AutoRNC’s sampled conformations according to, for example, their electrostatic and hydrophobic interactions. Given that we have shown here that AutoRNC already produces (equally weighted) conformations that nicely reproduce the conformational properties of proteins in their unfolded states, care would be needed to ensure that this useful feature is not destroyed by any added post-processing scheme.

## Methods

### Using AutoRNC

The release version of AutoRNC contains a number of utilities and scripts, but the main source code operates as a single executable program that should be straightforward to compile on any modern computing system. The code is accompanied by pre-built libraries of dipeptide conformations sampled from the RCSB, and three pre-processed ribosome template structures that include the systems most widely used in recent experimental studies (see below). If regions of tertiary structure are desired to be built into AutoRNC’s models, the user is required to provide a structure of the complete nascent chain construct (see below): the code will retain those residues that are part of specified tertiary structure elements and will discard and rebuild those residues that are specified by the user as being unfolded.

### Dipeptide library construction

A central assumption underpinning AutoRNC is that the conformational properties of unfolded regions of nascent chains can be reproduced by building chains via the superposition of dipeptide fragments extracted from the RCSB. To this end, we constructed a number of dipeptide libraries as follows. The Dunbrack group’s PISCES webserver (41) was used to select a non-redundant set of protein structures solved at resolutions better than 1.6 Å with maximum R-values of 0.25 or better and with less than 50% identity between the chains; at the time of download, the resulting dataset contained 6791 protein chains. Each chain was processed to remove any hydrogen atoms, and to renumber residues where necessary, before being run through the secondary structure assignment program DSSP developed by Kabsch and Sander (42). The resulting assignments were then used to extract dipeptides from within each chain and add them to an appropriate library. Candidate dipeptides were checked to ensure that they satisfied a number of conditions: all sidechain atoms must be present, the occupancies for all atoms in both residues must be 1, the residues must be adjacent in sequence, and the distance between their Cα atoms must be < 4.0 Å; finally, dipeptides involving either the first or the last residue in each chain were ignored.

Given that there are twenty common amino acids, there are 400 possible dipeptide sequences. For each possible sequence, we assembled seven libraries, each representing a different combination of the secondary structures assigned to the N-terminal and C-terminal residues of each dipeptide. The libraries cover the following combinations: (1) coil+coil, (2) helix+helix, (3) sheet+sheet, (4) helix+other, (5) other+helix, (6) sheet+other (7) other+sheet; here, “helix+other”, for example indicates that the N-terminal residue of the dipeptide is in an α-helical conformation while the C-terminal residue is in any non-helical conformation. In assigning conformations to the appropriate library, we interpreted DSSP’s alphabet in the following way: residues assigned a letter ‘H’ were treated as α-helical, those assigned a letter ‘E’ were treated as β-sheet, and any residue that was not assigned a letter ‘H’, ‘E’, ‘B’ (β-bridge), or ‘G’ (3,10-helix) was treated as a coil residue. The first of these seven libraries is used for all model-building of unfolded regions; the remaining libraries are used only when the user specifies elements of secondary structure that are to be built into the models. For example, to build an α-helix embedded in the hypothetical sequence `GLATLQAN` at residues 3-7, AutoRNC would make use of the following dipeptide libraries: `GL`(`CC`), `LA`(`OH`), `AT`(`HH`), `TL`(`HH`), `LQ`(`HH`), `QA`(`HO`), `AN`(`CC`) where `CC` represents “coil+coil”, `OH` represents “other+helix”, `HH` represents “helix+helix”, etc. The dipeptide libraries constructed above contain substantial numbers of conformations to sample from: for example, the median number of conformations present in the 400 “coil+coil” libraries is 828, with the minimum number in any such library being 50 (for the dipeptide MC) and the maximum number being 5897 (for the dipeptide GG).

### Ribosome template structure selection

In principle, any ribosome structure can be used with AutoRNC provided that it contains at least one amino acid residue attached to the tRNA in the PTC (see below). Prior to being used in AutoRNC a modest amount of pre-processing of the ribosome structure is required; this takes the form mainly of renaming chains and `ATOM` entries so that AutoRNC successfully recognizes the nascent chain from the other protein chains of the ribosome. The three pre-processed ribosome structures that are provided as templates with the AutoRNC are the following: (1) RCSB ID 5UYM (43), which we think serves as a good model for an elongating ribosome and for RNC constructs that have been generated experimentally either by flash-freezing or by tRNA starvation methods; (2) RCSB ID 3JBV (44), which we use for RNC constructs that contain a SecM arrest sequence at their C-terminus; and (3) RCSB ID 7OIZ (45), which we use for RNC constructs that contain a TnaC arrest sequence. Reflecting the frequency with which RNC constructs are designed in the literature, the vast majority of the RNC models built here make use of templates 5UYM and 3JBV.

### Generating conformational models with AutoRNC

AutoRNC builds models starting with the C-terminal residue of the construct and working forwards toward the N-terminus. In its usual mode of operation, the C-terminal residue will be the amino acid directly attached to the PTC tRNA (in the case of the 5UYM-based template), or the N-terminal-most residue of the arrest sequence (in the case of the 3JBV or 7OIZ-based templates). Further residues are added to the nascent chain one at a time, by superimposing dipeptide conformations onto the pre-existing chain using atoms of the C-terminal residue of the dipeptide and retaining the N-terminal residue of the dipeptide if the superposition satisfies selection criteria (see below). If the new residue to be added is part of a user-specified tertiary element, then the dipeptide conformation is extracted directly from the structure provided by the user; if the new residue is instead part of a user-specified secondary structure element, then the dipeptide conformation is selected randomly from the pre-made dipeptide library that has the appropriate sequence and the appropriate backbone conformation (see above); finally, if the new residue is not part of a user-specified secondary or tertiary structure element, then the dipeptide conformation is selected randomly from the pre-made “coil+coil” library with the appropriate sequence.

Prior to accepting the new residue, the following selection criteria are applied. First, the root mean square deviation (RMSD) of the superimposed atoms (Cα, N, C, O) is measured, and the conformation is rejected if the RMSD exceeds a user-specified threshold; in the applications described here, this threshold was generally set to a relatively permissive value of 0.50 Å, but more stringent values (e.g. 0.25 Å) can often be used while still generating models relatively rapidly. If the RMSD criterion is satisfied, then the newly added amino acid is checked for steric clashes with the rest of the nascent chain and with the ribosome, with different clash thresholds definable for the two categories. Intra-nascent chain clashes are monitored between the newly added residue and all previously placed atoms of the nascent chain other than those of the residue to which it is peptide-bonded; in most of the constructs reported here, intra-chain pairs of atoms were considered to be clashing if their distance was < 3.0 Å. Inter-chain clashes are monitored between the newly added residue and all atoms of the ribosome; in most of the constructs reported here, pairs of atoms were considered to be clashing if their distance was < 3.5 Å. To accelerate the acceptance of models AutoRNC allows users to relax these distance thresholds for residues closest to the C-terminus of the nascent chain; in most of the constructs reported here, the distance thresholds applied to the last ten residues were scaled upwards by 33%.

If models are desired to be generated more rapidly it is possible to relax the effective clash criteria further in a variety of ways. One possibility is to alter the desired resolution of the nascent chain models: in addition to allowing full atomic models to be generated, AutoRNC allows users to request models that contain only backbone atoms, or backbone atoms + Cβ atoms; for obvious reasons, backbone-only models are much less prone to being involved in steric clashes. A second possibility for more rapidly obtaining models is to retain the atomic representation of the models while decreasing the two distance thresholds used to determine clashes for all residues of the nascent chain. Rough guidelines for deriving relaxed distance criteria can be obtained from examination of the closest inter-atomic distances in the cryo-electron microscopy structures used as ribosome templates. For example, the smallest distances measured between two non-bonded atoms in the nascent chains of those ribosome templates that contain a nascent chain are 2.1 Å for 3JBV and 2.2 Å for 7OIZ; similarly, the smallest distances measured between atoms in the nascent chain and atoms of the ribosome (excluding the PTC tRNA) are 2.1 Å for 3JBV and 2.4 Å for 7OIZ. These results suggest that threshold distances as low as 2.0 Å might reasonably be used to identify both intra-nascent chain and nascent chain-ribosome clashes.

If no steric clashes are present after the superposition, the new residue is accepted and AutoRNC proceeds to the next unmodeled residue. If clashes are present, then the superimposed residue is rejected, and one of the following scenarios plays out. If the residue to be added is part of an unfolded region of the protein or is part of a secondary structure element defined by the user, then additional conformations are selected from the appropriate dipeptide library and checked for acceptance in the manner described above; this continues until all possible conformations in the library have been tried or until the total number of failures reaches a user-specified threshold which, for the models presented here, was set to 250. If, on the other hand, the residue is part of a tertiary structure element defined by the user, then there is (currently) no alternative conformation to sample, and the failure threshold is immediately assumed to have been reached. One possibility when reaching the failure threshold is to abandon the current model of the nascent chain entirely; doing so, however, can represent a significant loss in terms of the time invested in attempting to build the model. AutoRNC, therefore, allows an alternative of “backtracking” the model by a specified number of residues (N_back_) instead; when this occurs, the failure count is reset to zero. It is possible for repeated backtracking events to occur, with in some cases the nascent chain backtracking all the way to the final residue, but if the number of backtracking events exceeds a second threshold (set to 100 here) for a given model, the chain is restarted from scratch. In most of the constructs built here, the backtracking depth was set to 5, but for constructs containing secondary structure elements it was set to one residue larger than the largest secondary structure element: backtracking by this depth ensures that AutoRNC will backtrack to a residue that is C-terminal to the secondary structure element, thereby allowing it to sample different orientations of the element and preventing it from otherwise continually attempting to lengthen a secondary structure element that has no prospect of fitting within the ribosome tunnel.

Once all residues of the nascent chain have been successfully built for a given model, it is immediately written to an output PDB file, and the process restarts until the desired number of models are built. A video example of the entire process is shown in Movie S1.

### Modeling the conformational properties of unfolded proteins

To demonstrate that AutoRNC’s chain-building routines produce reasonable conformational distributions for the unfolded regions of proteins in the absence of a ribosome – something for which methods are already available (e.g. (46)) – AutoRNC was used to build structural ensembles for the following four model proteins: protein L, ubiquitin, acylphosphatase, and the N-terminal domain of pertactin. These four proteins were selected owing to the availability of experimental radii of gyration, R_gyr_, for their unfolded states in native conditions (reviewed in (30)). 10,000 different models of each protein were generated (in the absence of any ribosome) using AutoRNC’s default input parameters (see above), with each such model being built starting from the C-terminus with a different conformation randomly selected from the appropriate dipeptide library. The R_gyr_ value of each model was calculated using an in-house script and an overall mean value was obtained as the square root of the 10,000 mean squared values. To demonstrate that the directionality with which AutoRNC builds polypeptide chains is largely immaterial, we built a further 10,000 models starting from the N-terminus. Interestingly, the computed mean R_gyr_ values for models built from the N-terminus were systematically slightly higher than those built from the C-terminus; however, the maximum difference found for any of the four proteins was only 4 %.

To verify that the conformational properties of unfolded regions are appropriately sampled by AutoRNC, a random 1000-amino acid sequence was generated using the Sequence Manipulation Suite (47) and 1000 models were again constructed with the same protocol. The phi and psi angles of all non-gly, non-pro, and non-pre-pro residues in all 1000 models were measured using an in-house script and the resulting 2D histogram was visualized using R (RStudio Team, 2020).

### Collection of RNCs whose folding status has been determined experimentally

To demonstrate the capabilities of AutoRNC in its intended setting, we surveyed the literature for RNC constructs that contained varying degrees of structure in their nascent chains. In total, we identified ∼60 RNCs from the literature that include experimental evidence suggesting which residues in the nascent chain, if any, should be modeled as folded. The total number of constructs considered in these papers far exceeds 60, but since many of these vary by only point mutations, we instead chose RNCs that covered the various degrees of nascent chain folding shown to occur in each of these works. In general, we found that all RNCs could be roughly assigned to one of the five categories illustrated in Figure 2. When not explicitly provided, we determined the sequence of each construct from source information provided in the paper: e.g., plasmid sequences.

To demonstrate AutoRNC’s ability to build elements of tertiary structure into its models, we created folded template structures of all full-length constructs (including any uncleaved tag residues) using a local installation of ColabFold (28) that incorporates the AlphaFold2 neural networks (27) for protein structure predictions. Specifically, we predicted 5 models with 20 recycles and used the model with the best predicted TM-score as the template structure for the fully folded nascent chain. We note that, for the constructs studied here, all domains that have been shown to be folded have plausible template structures already available in the RCSB: Table S1, for example, identifies the best available match identified for each construct by a BLAST search (48) of the SEQATOMs database (49). However, using AlphaFold2-predicted structures in place of these available structures allows us to: (1) use a consistent protocol for all RNCs, (2) include residues that are missing from the solved structures, and (3) model chimeric constructs that contain structured domains from different proteins.

All AutoRNC runs were first performed with the parameter settings identified above. In those cases where AutoRNC appeared to have difficulty returning models rapidly, the alternative strategies outlined above were used. For some RNCs, it was possible to obtain models by switching the resolution of the desired models from fully atomic to backbone only; for others, it was possible to obtain models by retaining full atomic resolution but dropping the distance thresholds used to identify steric clashes to 2.0 Å as suggested above. Finally, even with these adjustments made, in two particularly recalcitrant cases (see main text), a hairpin loop in protein L24 needed to be removed before two constructs containing folded domains could be successfully built. Table S1 identifies those cases where AutoRNC’s input parameters were adjusted during model building.

## Code availability

The AutoRNC source code, together with the libraries of dipeptide conformations used to build unfolded regions of nascent proteins, and all of the RNC models reported here, will be made available for download at the following GitHub repository: https://github.com/Elcock-Lab/AutoRNC/.

## Acknowledgements

This research was supported in part through computational resources provided by The University of Iowa, Iowa City, Iowa.

