## Supporting Information for "AutoRNC: an automated modeling program for building atomic models of ribosome-nascent chain complexes"

### Figure Legends

**Figure S1** **AutoRNC accurately models unfolded protein conformations.** (A) Histograms of the radius of gyration, $R_{gyr}$, of 10,000 conformations of four proteins sampled in their unfolded states using AutoRNC with default input parameters. (B) Distribution of Ramachandran angles in 1000 conformations of a 1000-amino acid protein with a randomly generated sequence. Only the Ramachandran angles of the non-glycine, non-proline, and non-pre-proline residues are shown.

**Figure S2** **Gallery of experimental constructs built using AutoRNC with an elongating ribosome template.** Each image displays a single RNC construct built using secondary and tertiary structure elements taken from experimental sources (see Table S1 for references). Ribosomes are shown as transparent cartoons colored by chain, the PTC tRNA chains are shown as opaque cartoons and modeled nascent chains are shown as opaque blue cartoons. Constructs are numbered according to the numbering scheme used in Table S1. All models in this figure were built using the 5UYM ribosome template structure.

**Figure S3** **Gallery of experimental constructs built using AutoRNC with a ribosome templates containing stall sequences.** Same as Figure S2 but showing RNC constructs that contained either SecM (models 31-57) or TnaC (models 58 & 59) stall sequences at their C-termini.

**Figure S5** **AutoRNC models can reflect idiosyncrasies of the ribosome templates.** (A) In the ribosome template structure that was stalled via the *E. coli* TnaC arrest peptide (7OIZ), the hairpin loop of ribosomal protein L24 partially occludes the exit tunnel; this can lead to clashes with RNCs known to fold (see main text). (B) 1000 RNC models of a 441-residue EF-G construct built using the non-stalled ribosome template structure 5UYM. Only the last 60 residues are displayed (opaque cyan cartoon) in order to highlight the three distinct exit tunnels that are identified. (C) The ribosome template structure stalled with the *E. coli* SecM arrest peptide (3JBV) contains large gaps between the tRNA binding sites. In cases where AutoRNC has difficulty building non-clashing models that use the more conventional exit tunnels, it can occasionally build models in which the nascent chain "doubles back" to occupy these regions.

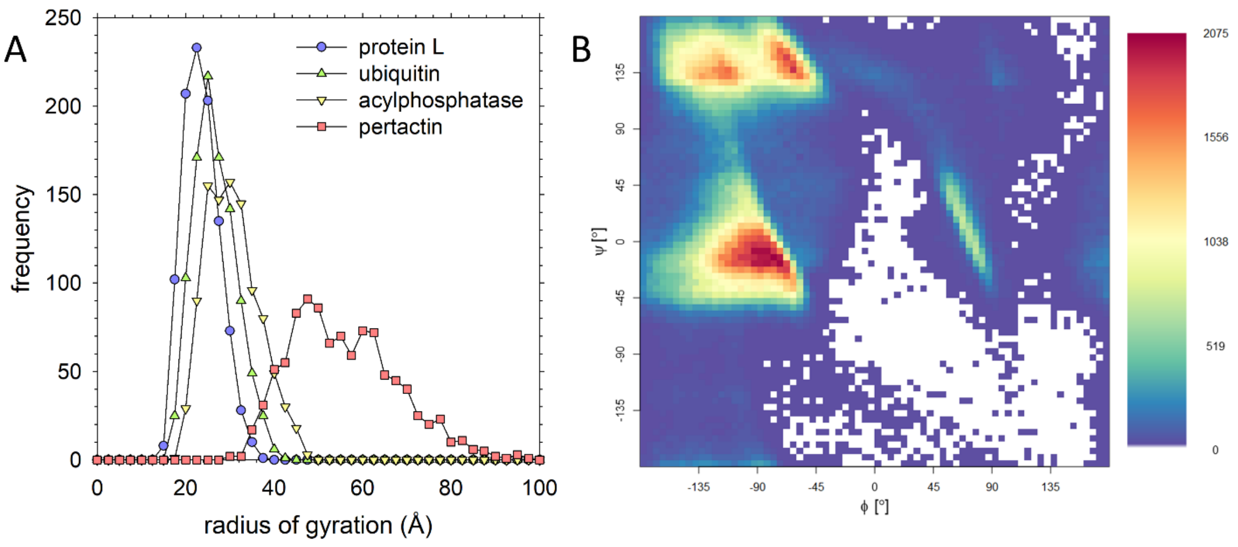
A
B
protein L
ubiquitin
acylphosphatase
pertactin
frequency
radius of gyration (Å)
ψ [°]
ϕ [°]
2075
1556
1038
519
0

**Figure S1**

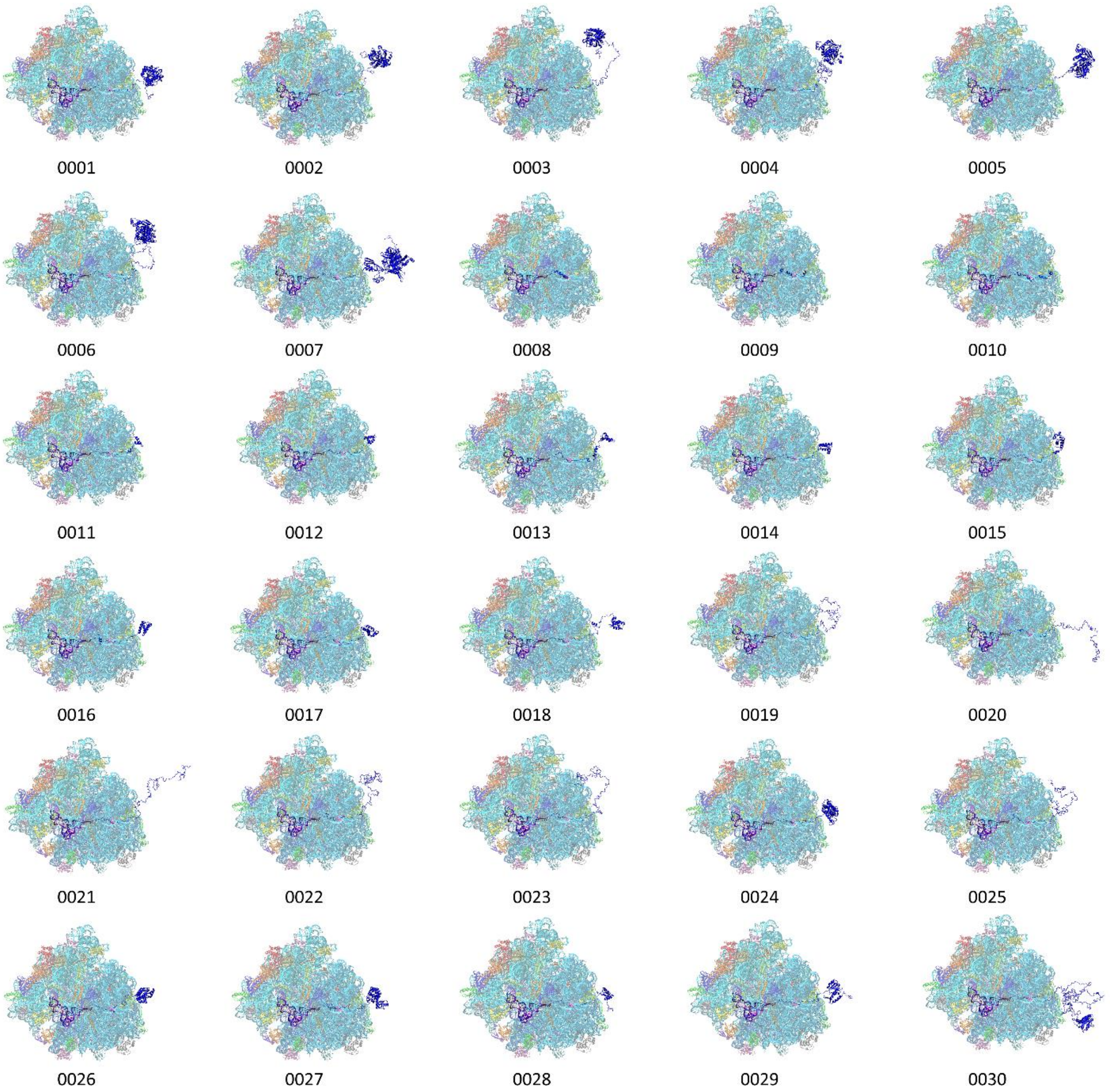

**Figure S2**

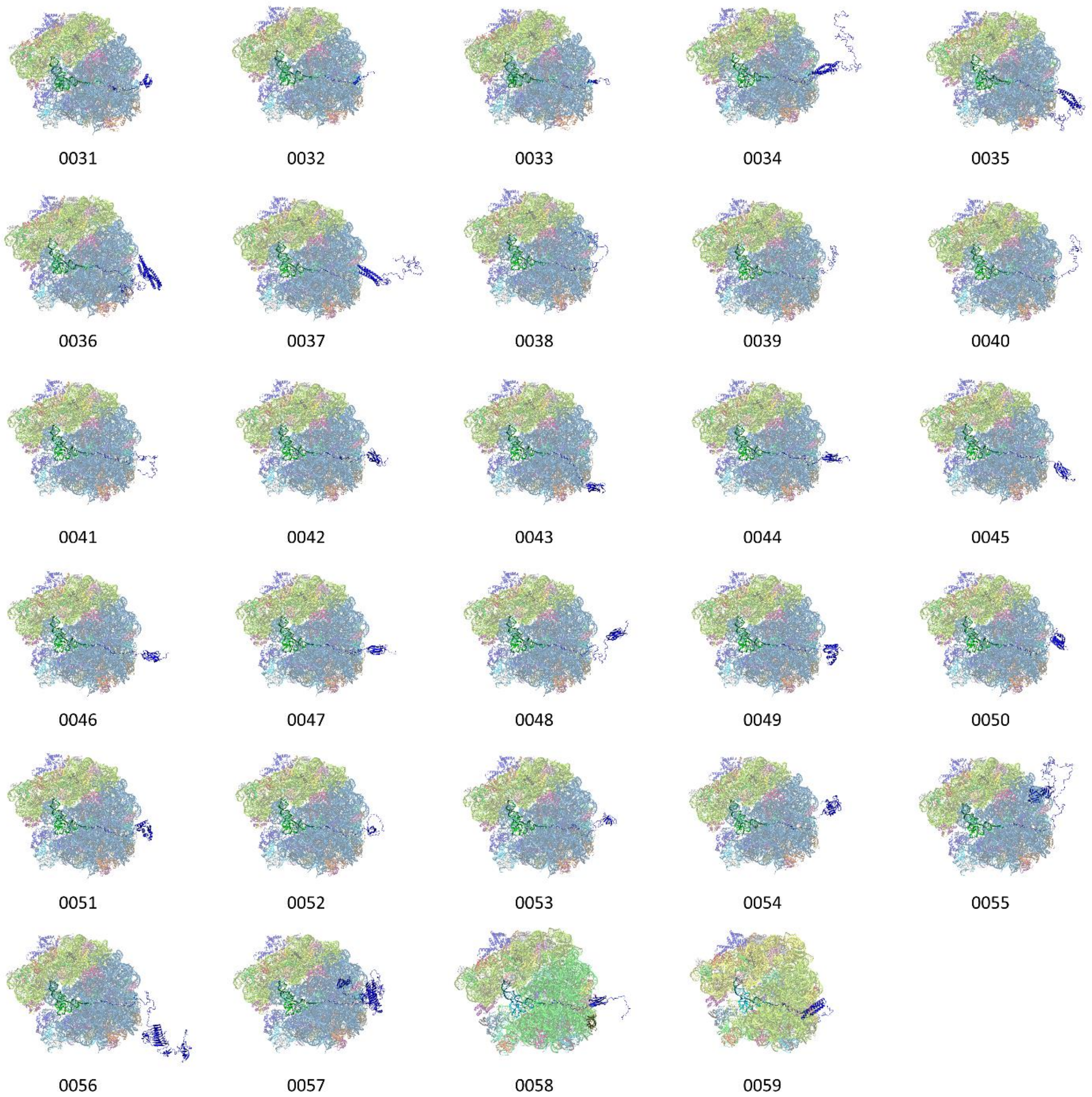

**Figure S3**

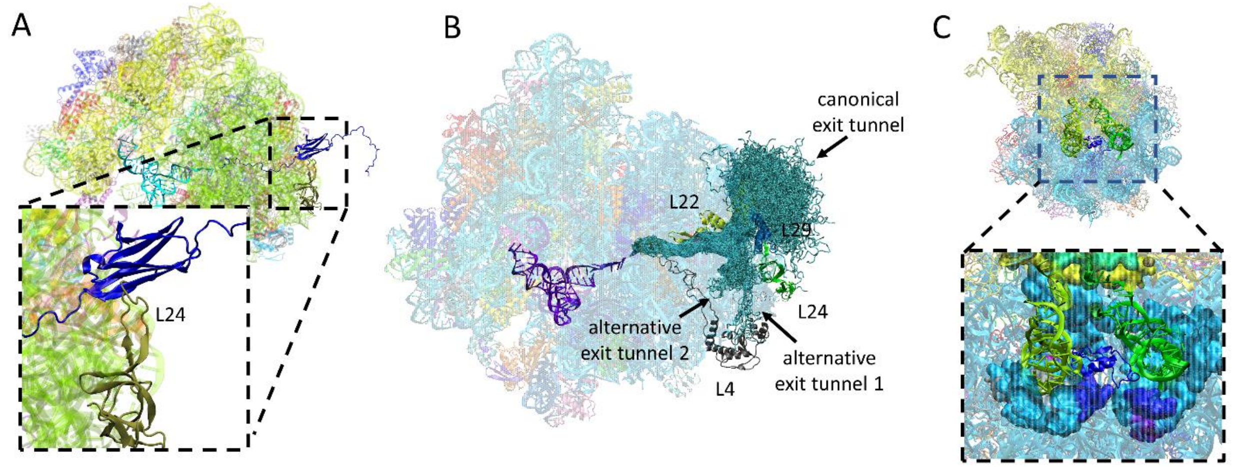
A
L24
B
canonical
exit tunnel
L22
L29
L24
alternative
exit tunnel 2
alternative
exit tunnel 1
L4
C

**Figure S4**
